# HIRA-dependent provision of histone H3.3 in active chromatin ensures genome compartmentalisation

**DOI:** 10.1101/2024.08.27.609896

**Authors:** T. Karagyozova, A. Gatto, A. Forest, J.-P. Quivy, M. Marti-Renom, L. Mirny, G. Almouzni

## Abstract

The mammalian genome, organised into chromatin, adopts a three-dimensional (3D) folding within the cell nucleus with spatially segregated active and repressed compartments, termed A and B. However, how nucleosome deposition impacts these levels of organisation is unknown. Here, we monitored changes in 3D genome folding by Hi-C after impairing the chaperone HIRA, involved in histone H3.3 deposition. In the absence of HIRA, H3.3 enrichment decreases in compartment A that also shows weaker interactions. At this scale, histone post-translational modifications (PTMs) do not follow H3.3 changes. In line with impaired H3.3 nucleosome maintenance, compartment A accessibility measured by ATAC-seq increases. Specifically, at active genes, accessibility increases in gene bodies but decreases at promoters where compensation by H3.1 reduces nucleosome turnover. Notably, regions flanking active genes show reduced insulation. We conclude that the HIRA-dependent pathway involved in H3.3 deposition is key to maintain higher order organisation in active regions and impact compartmentalisation independently of histone PTMs.

## Introduction

The organisation of chromatin ranges from a basic unit, the nucleosome up to higher order level folding within the nucleus. The nucleosome comprises ∼146bp DNA wrapped around an octamer of four core histones, H2A, H2B, H3 and H4 (Luger et al., 1997) and linker DNA (Van Holde, 1989), a repeated module that forms the nucleofilament. The incorporation of different histone variants and their post-translational modifications (PTMs) provide versatility in the basic module. Beyond the nucleofilament, folding occurs with loops (Rao et al., 2014) and topologically-associating domains (TADs), each enriched in self-interactions and insulated from neighbouring regions (Davidson and Peters, 2021; Dixon et al., 2012; Nora et al., 2012). A further level of higher organization is the partitioning into compartments A and B, which are spatially segregated from each other and correspond respectively to open, active euchromatin and dense, repressive heterochromatin (Jerkovic and Cavalli, 2021; Lieberman-Aiden et al., 2009; Magnitov and De Wit, 2024). Surprisingly, while we have learnt a lot about loop extrusion as mechanism contributing to the dynamics of loops (Corsi et al., 2023; Fudenberg et al., 2016), the nucleosomal dimension with the assembly pathways and choices of histone variants has not been explored for its impact on genome folding in mammals.

Nucleosome formation relies on the use of distinct histone variants escorted by histone chaperones operating in a controlled manner in space and time (Delaney et al., 2023). Indeed, two distinct mechanisms operate for histone H3-H4 deposition, a DNA synthesis-coupled (DSC) or independent (DSI) pathways. In S phase, the DSC pathway involving the CAF-1 complex (Smith and Stillman, 1989) deposits the replicative H3.1/2 (Tagami et al., 2004). This major provision of replicative H3 is critical in cycling cells (Franklin and Zweidler, 1977; Zweidler, 1980). In contrast, the major non-replicative H3 variant, H3.3, expressed throughout the cell cycle (Wu and Bonner, 1981) provides another source and is incorporated independently of DNA synthesis at any time (Ahmad and Henikoff, 2002). The chaperone complex HIRA is key for this pathway (Ray-Gallet et al., 2002) to promote H3.3 deposition (Tagami et al., 2004). H3.3 enrichment at active regions and regulatory elements (Goldberg et al., 2010) depends on HIRA, possibly due to its interaction with RNA Pol II (Ray-Gallet et al., 2011) or transcription factors (Pchelintsev et al., 2013). This H3.3 genomic distribution is paralleled by active PTM patterns (Goldberg et al., 2010; Mito et al., 2005). In addition, independently of transcription, HIRA can promote H3.3 incorporation at nucleosome gaps, potentially through its ability to bind naked DNA (Ray-Gallet et al., 2011). Finally, enrichment of H3.3 has also been reported at repetitive elements and depends on ATRX/DAXX (Drané et al., 2010; Lewis et al., 2010). While we recently learnt that the general histone chaperone FACT impacts nucleosome occupancy and active gene organisation in human cells (Žumer et al., 2024), we still do not know whether and how HIRA-mediated H3.3 incorporation at active chromatin impacts its organisation in 3D. This is even more important considering that HIRA proved important for both *de novo* deposition and recycling of H3.3 in the context of transcription (Torné et al., 2020). This is particularly intriguing considering early mammalian development, when higher-order chromatin organisation is being established (Du et al., 2017; Gassler et al., 2017; Ke et al., 2017) concomitantly with a genome-wide H3.3 redistribution (Ishiuchi et al., 2020). Therefore, we hypothesized that HIRA-mediated nucleosome assembly could also have an important function in active chromatin organisation.

In this work, we explored how perturbing chromatin assembly by impairing the H3.3-specific chaperone HIRA could impact the different scales of higher-order genome organisation. We found that HIRA ensures enrichment of H3.3 within compartment A. Indeed, in the absence of HIRA, H3.3 enrichment decreases significantly in compartment A, but this occurs without major compartment A to compartment B (A-to-B) switching. This decrease is accompanied by reduced compartment A interactions in cells lacking HIRA. Strikingly, the absence of HIRA did not lead to a similar redistribution of H3 PTMs at the scale of compartments. However, accessibility in compartment A increased as measured by ATAC-seq. Further analysis of the ATAC-seq and Hi-C data at higher resolution showed that accessibility of active gene bodies increased, and local folding around these genes was perturbed in the absence of HIRA. We conclude that efficient H3.3 nucleosome formation in active chromatin contributes to its compartmentalisation independently of histone PTMs.

## Results

### HIRA ensures proper provision of H3.3 in compartment A

To investigate the impact of distinct histone variant deposition on higher-order chromatin organisation, we focused on the H3.3-specific chaperone HIRA. We used a previously characterized constitutive knock-out (KO) of HIRA in HeLa cells (Ray-Gallet et al., 2018). We compared 3D genome organization in wild-type (WT) or HIRA KO cells by Hi-C. In parallel, we assessed the distribution of H3.3, H3.1, a set of H3 PTMs and chromatin accessibility (Figure 1A). To avoid S phase heterogeneity due to their distinct incorporation (Delaney et al., 2023), we used the ChIP-seq data from G1/S cells (Gatto et al., 2022) to profile H3 variant enrichment. We first verified that Hi-C maps from the two parental H3.1-SNAP and H3.3-SNAP cell lines showed comparable *cis*/*trans* contacts (Supplementary Figure 1A) and compartment calls (based on eigenvector (EV) decomposition (Lieberman-Aiden et al., 2009) at 50kb resolution, Supplementary Figure 1B). We found a strong enrichment of H3.3 in compartment A and relative depletion in B (Figure 1B). In contrast, H3.1 was rather depleted in A and enriched only in large (>2Mb) compartment B domains (Figure 1B). We then carried out Hi-C analysis in HIRA KO. The maps obtained from HIRA KO resembled those of WT cells (Figure 1B, C, left) based on their *cis* contact proportion (Supplementary Figure 1A) and contact distance decay (Supplementary Figure 1C). The proportion of compartment changes of 2.1% A-to-B and 1% B-to-A (Figure 1D) remains limited yet exceeds variability between the cell lines (Supplementary Figure 1B). However, in the absence of HIRA, H3.3 enrichment decreased throughout compartment A (Figure 1B, C, quantified in Supplementary Figure 1D). In contrast, H3.1 did not show substantial changes (Figure 1B, C, quantified Supplementary Figure 1D). Finally, at compartment borders we observed a less sharp transition for both variants, which is reminiscent of previously reported blurring at H3.3-enriched sites (Gatto et al., 2022). As represented schematically (Figure 1E), we conclude that lack of HIRA leads to a general decrease of H3.3 enrichment in compartment A without a strong impact on compartment identity.

**Figure 1.**
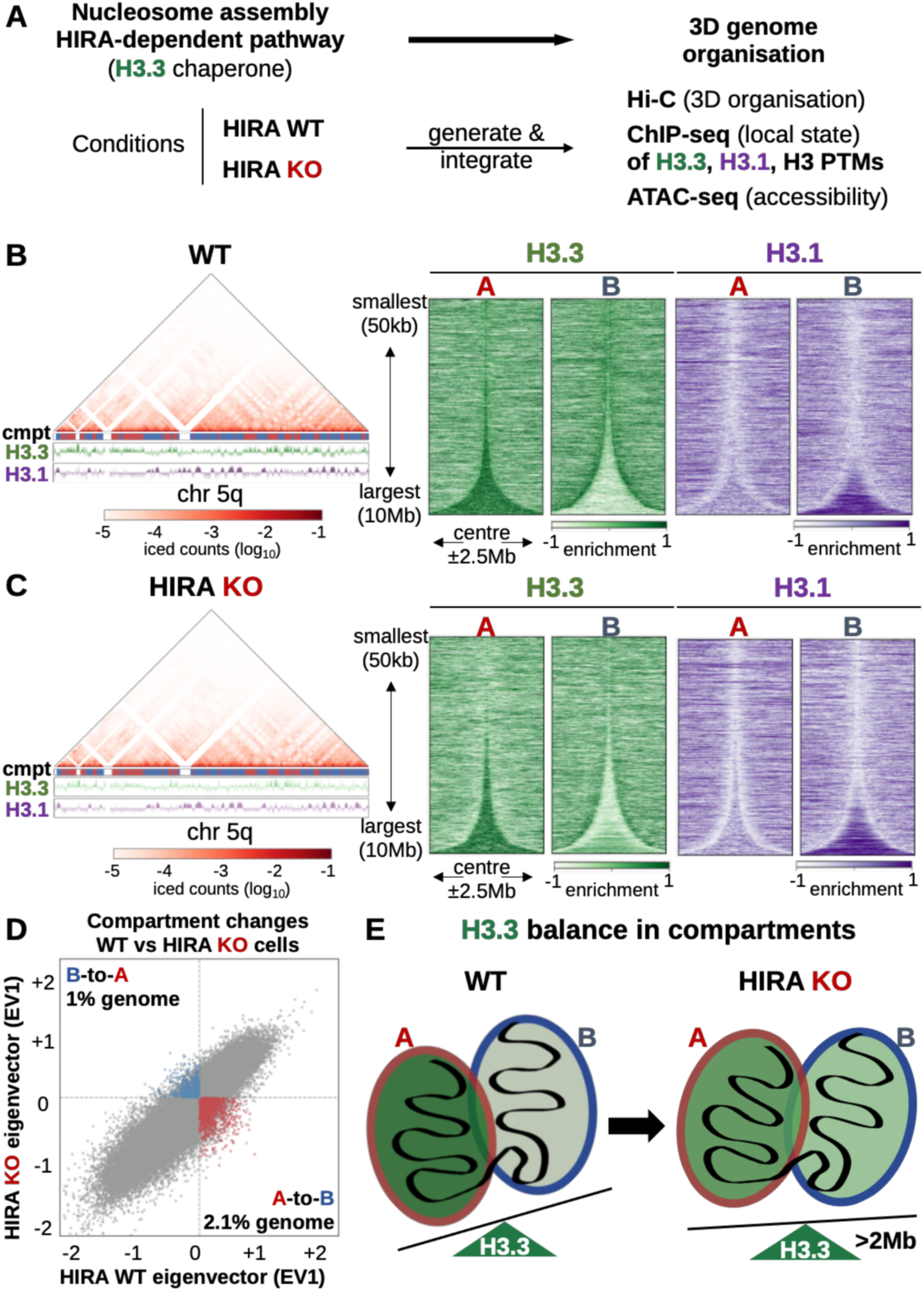
HIRA ensures proper provision of H3.3 in compartment A. **A.** Experimental strategy to assess the effect of disrupting chromatin assembly on higher-order genome organization by constitutive HIRA knock-out (KO). Hi-C was performed in WT and HIRA KO H3.1-SNAP and H3.3-SNAP cells. To compare the 3D folding of chromatin to local state, we obtained H3.1 and H3.3 SNAP ChIP-seq from Gatto et al. (2022) and profiled H3 PTMs by ChIP-seq and accessibility by ATAC-seq. **B.** Left: Representative Hi-C map at 50kb resolution with A/B compartment track and H3.1 and H3.3 enrichment at 10kb resolution of chromosome 5q (chr5: 50-170Mb) from WT cells. Right: Total H3.3 and H3.1 enrichment at 10kb bins at A (n = 1573) and B (n = 1573) compartment domains from WT HeLa cells, sorted by size and centered at their middle ±2.5Mb. **C.** Left: Hi-C map, compartment track and right: H3.1 and H3.3 enrichment shown in A (n = 1582) and B (n = 1588) compartment domains as in **B.** for HIRA KO cells. **D.** EV1 (1^st^ eigenvector, indicating compartment) of 50kb-binned Hi-C matrices from HIRA WT vs KO cells. Bins which change from A-to-B (lower right quadrant) or B-to-A (upper left quadrant) in the same direction in both cell lines are coloured red and blue, respectively). **E.** Schematic of H3.3 balance in compartments A and B in WT and HIRA KO cells, showing reduced H3.3 enrichment in compartment A and increased detection of H3.3 in large (>2Mb) compartment B domains. Enrichment shown is z-score of log_2_ IP/input.

### Compartment A decreases its interactions in the absence of HIRA without TAD reorganisation

Given the observed changes in H3.3 enrichment in compartment A in HIRA KO cells, we wondered if this affected the strength of interactions within compartments. Differential maps revealed reduced contact frequency between A-A and A-B in parallel with increased interactions between B-B regions in *cis* in HIRA KO (Figure 2A). Saddle plot analysis confirmed this observation genome-wide (Figure 2B (red box), 2C) in both cell lines (Supplementary Figure 1E, F). Additionally, this analysis also revealed that B interacted more with intermediate (I) compartments which were neither A nor B (Figure 2B (blue box), 2C, Supplementary Figure 1E, F). Given these observations at compartments, we next examined effects on chromatin organization at finer scales. We identified TADs from HIRA WT and KO cells at 10kb resolution using the insulation score metric (Crane et al., 2015). TAD borders overlapped well between the two conditions (Supplementary Table 1) to the same extent as H3.1- and H3.3-SNAP cell lines. Since the variants displayed compartment-specific enrichment (Figure 1B, C, Supplementary Figure 1D), we further profiled their patterns at TAD borders within compartment A or B individually (Figure 2D). In WT cells, while H3.3 peaked at TAD borders in both A and B compartment, consistent with gene enrichment there (Dixon et al., 2012), H3.1 was depleted in compartment B (Figure 2D). In the absence of HIRA, at TAD borders H3.3 became blurred in compartment A and lost in compartment B, while the H3.1 depletion in compartment B weakened. For a higher resolution analysis, we focused on CTCF sites, typically enriched at TAD borders (Dixon et al., 2012). In WT cells, H3.3 flanked CTCF sites, whereas in HIRA KO, H3.3 enrichment was reduced and peaked at the site (Supplementary Figure 2A). However, in both HIRA WT and KO cells the local insulation at TAD borders and CTCF sites (Figure 2E, Supplementary Figure 2B) remained comparable. Thus, HIRA proved important for maintaining A compartment interactions genome-wide and restricting interactions of B compartment to itself without affecting TAD-scale organisation.

**Figure 2.**
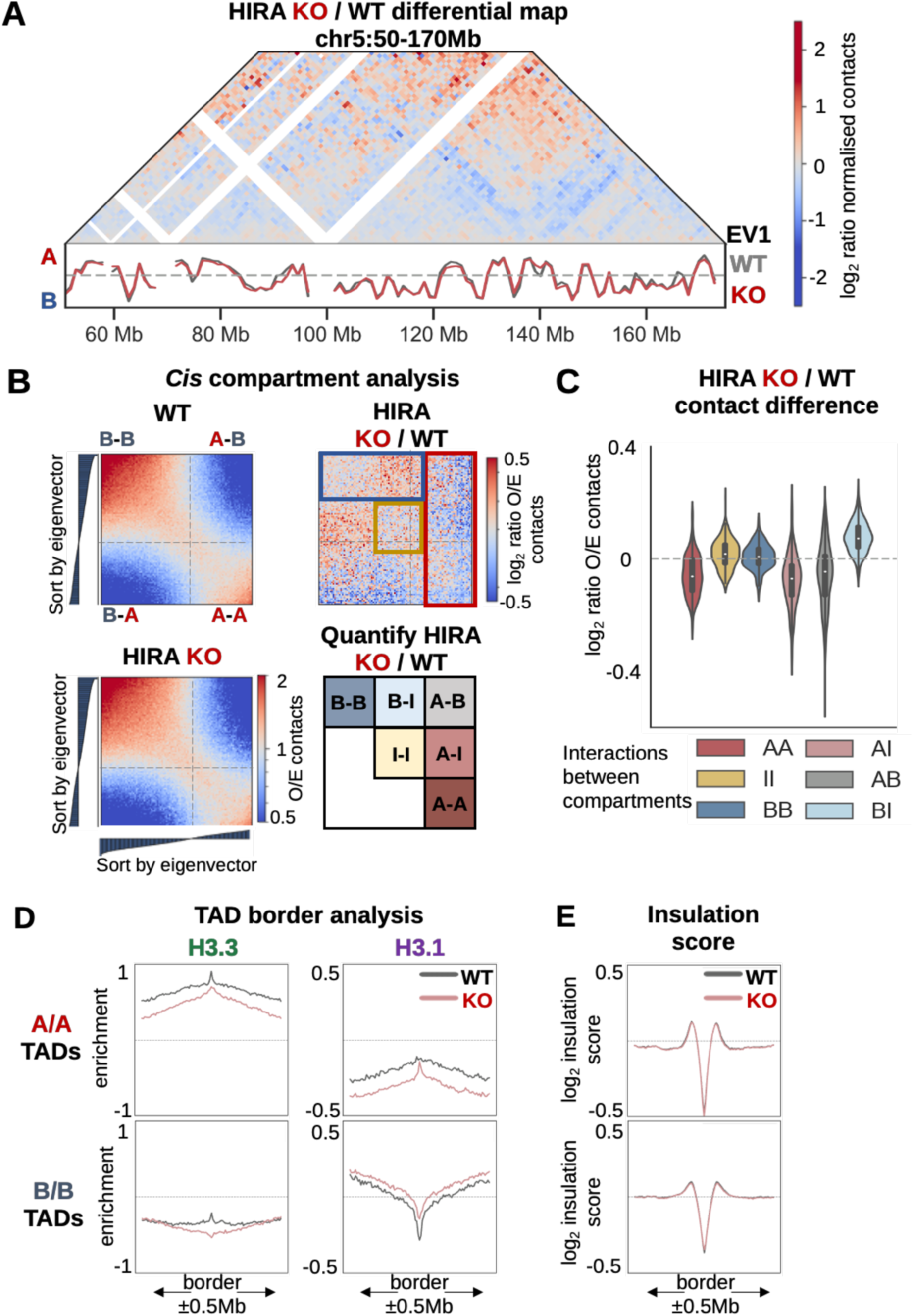
Compartment A domains decrease their interactions in the absence of HIRA without TAD reorganisation A. Differential Hi-C map (log_2_ ratio of normalised counts from HIRA KO/WT) at 1Mb resolution of chromosome 5:50-170Mb from 0-50Mb away from the diagonal. Tracks below show EV1 (HIRA WT in grey, KO in red). B. Left: Saddle plots from HIRA WT and KO (O/E contacts) at 50kb resolution based on EV1 percentiles. Right: Differential saddle plot (log_2_ ratio of O/E contacts of HIRA KO/WT). Red quadrangle denotes decreased contacts of compartment A genome-wide, blue quadrangle denotes compartment B interactions with itself and intermediate (I) compartment, and yellow square denotes I-I contacts. Differences are quantified in Fig. 2C following the schematic representation of genome-wide interactions separated into A (active), B (inactive) and I (intermediate) compartments based on EV1 terciles shown below. C. Difference (log_2_ HIRA KO/WT ratio) of O/E contact frequency between the sets of A/B/I compartments based on differential saddle-plots from H3.1-SNAP cells (Fig. 2B). D. Mean H3.1 and H3.3 enrichment (z-score of log_2_ IP/input) at TAD borders within compartment A (n = 2969, top) and B (n = 2340, bottom) from WT and HIRA KO cells at 10kb bins. TADs are centered at their start border ±0.5Mb. E. Insulation score (log_2_, 10kb bins) at TAD borders within compartment A (top) and B (bottom) from H3.1-SNAP WT vs HIRA KO cells. TADs are centered at their start border ±0.5Mb.

### At the compartment scale, in the absence of HIRA, H3.3 redistribution is not accompanied by corresponding PTM changes

Given the reports showing that phosphorylation of the H3.3-specific S31 residue can promote H3K27ac deposition by p300 (Armache et al., 2020; Martire et al., 2019; Morozov et al., 2023) or inhibit H3K9me3 removal by KDM4B (Udugama et al., 2022), we investigated whether the redistribution of H3.3 in the absence of HIRA led to changes in H3 PTMs. We performed native ChIP-seq for a panel of selected active (H3K4me3, promoter-associated, H3K4me1, H3K27ac, enhancer-associated) and inactive (H3K9me3, constitutive heterochromatin and H3K27me3, facultative heterochromatin) H3 marks (Figure 1A). These analyses did not reveal changes of PTM patterns in A/B compartments that mirror the H3.3 redistribution (Figure 3, Supplementary Figure 3A). More specifically, where we observed an increase in H3.3 enrichment (Figure 1C, Supplementary Figure 1D), no gain of active or loss of inactive marks in large B compartments occurred (Figure 3A, Supplementary Figure 3A). We then examined closer PTMs at regulatory sites by calling peaks for H3K4me1, H3K4me3 and H3K27ac (Supplementary Table 2). While H3K4me3 did not change in WT versus KO Supplementary Figure 3B), a small decrease occurred for H3K4me1 and H3K27ac (Supplementary Figure 3C, D). However, in WT, H3.3 was enriched at all peaks and strongly decreased in HIRA KO, while H3.1 showed the opposite behaviour (Supplementary Figure 3B-D). Therefore, changes in the enrichment of H3.3 and the marks at PTM peaks do not correlate. In addition, the importance of HIRA for local enrichment of enhancer-associated PTMs is not translated to the compartment scale. Thus, we conclude that HIRA contributes to compartment organisation independently of H3 PTM enrichment.

**Figure 3.**
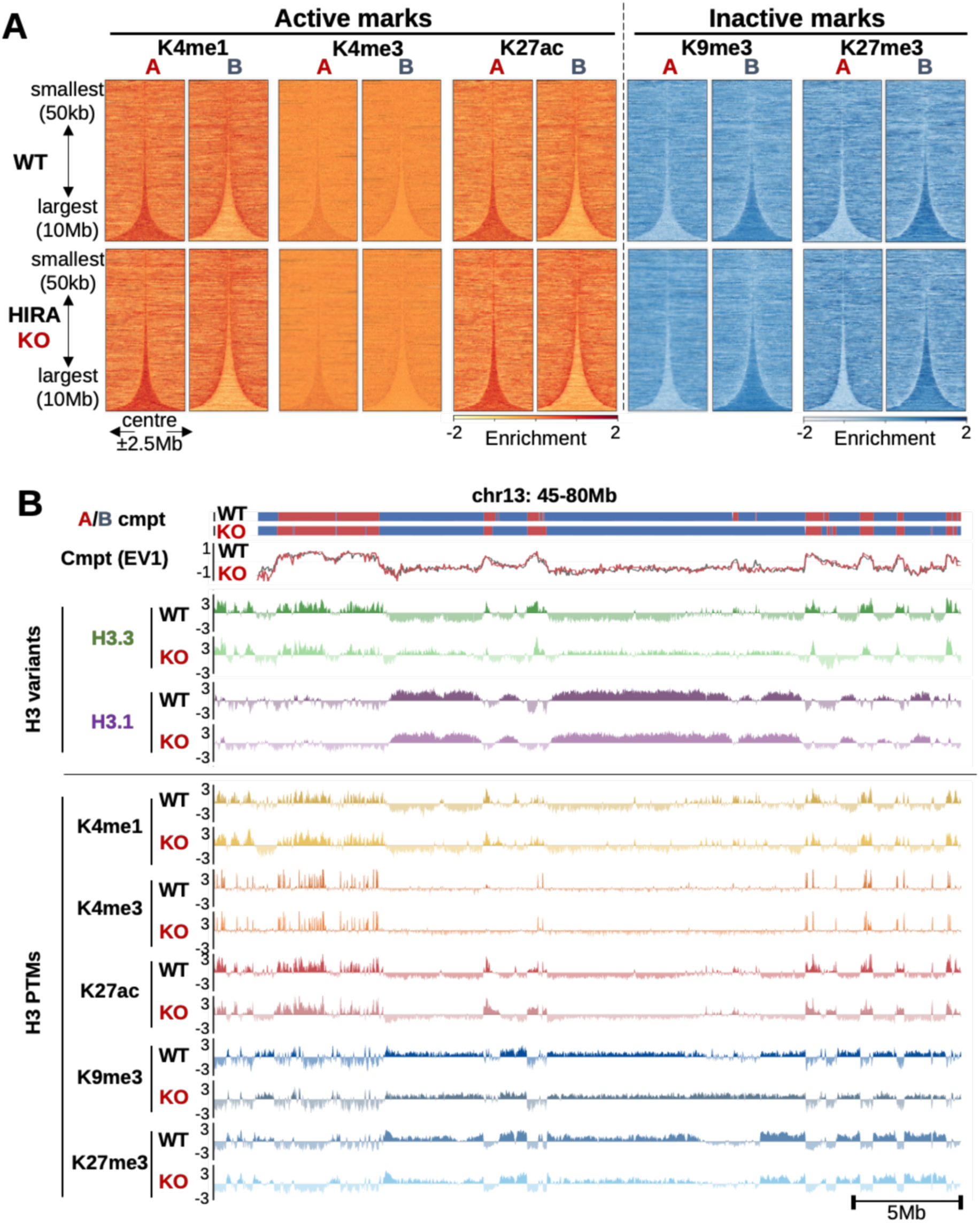
At the compartment scale, H3.3 redistribution in the absence of HIRA is not accompanied by corresponding H3 PTM changes. **A.** Active (H3K4me1/3, H3K27ac) and inactive (H3K9/27me3) PTM enrichment at 10kb bins at compartments A and B from WT and HIRA KO HeLa cells, sorted by size and centered at their middle ±2.5Mb. **B.** Compartment assignment and eigenvector tracks at 50kb resolution and enrichment of H3.3, H3.1, active (H3K4me1/3, H3K27ac) and repressive (H3K9/27me3) PTMs from HIRA WT and KO cells (chr13: 45-80Mb). ChIP-seq is shown at 10kb bins smoothed over 3 non-zero bins. H3.3, H3.1, PTM enrichment shown is z-score of log_2_ IP/input.

### Chromatin accessibility increases in compartment A and within active gene bodies in the absence of HIRA

Since transient HIRA depletion increased DNase I sensitivity (Ray-Gallet et al., 2011), we evaluated whether chromatin accessibility changes at the compartment level. We performed ATAC-seq in HIRA WT and KO cells (Figure 1A). We found that accessibility increased in compartment A and diminished in B in the absence of HIRA (Figure 4A). To investigate the defects at the nucleosomal scale, we called peaks from the ATAC-seq data, which overlapped well between the two conditions (Supplementary Table 3). In WT cells, ATAC-seq peaks showed an enrichment in H3.3 and a depletion of H3.1 (Supplementary Figure 4A). In the absence of HIRA, ATAC-seq and H3.3 decreased, while H3.1 enrichment increased at the peaks (Supplementary Figure 4A). Since this change at ATAC-seq peaks contrasted with the increased accessibility in compartment A, we then examined more broadly transcriptionally active regions, where HIRA plays a key role for deposition of H3.3 (Ray-Gallet et al., 2011; Schwartz and Ahmad, 2005; Torné et al., 2020). To examine accessibility over gene bodies, we classified genes based on their expression level from RNA-seq data and plotted total ATAC-seq signal (no fragment length filtering). In WT cells, active genes showed H3.3 enrichment, with only promoters highly accessible (Figure 4B, top). In HIRA KO, H3.3 enrichment and accessibility decreased concomitantly at active TSSs (Figure 4B, bottom), as observed at ATAC-seq peaks (Supplementary Figure 4A). Remarkably, the H3.3 decrease at active gene bodies paralleled an accessibility increase in a manner scaling with expression (Figure 4B). Notably, we did not detect substantial changes in pattern of gene expression globally (Supplementary Figure 4B) or in the top 5% expressed genes (only 2/879 downregulated in HIRA KO). We conclude that loss of HIRA leads to two distinct outcomes at active genes: (i) peaks at promoters lose both H3.3 and accessibility, (ii) at active gene bodies the broad loss of H3.3 is associated with increased accessibility. Given that gene bodies constitute a larger portion of compartment A regions compared to promoters, we propose that their increased accessibility is reflected at the level of compartment A.

**Figure 4.**
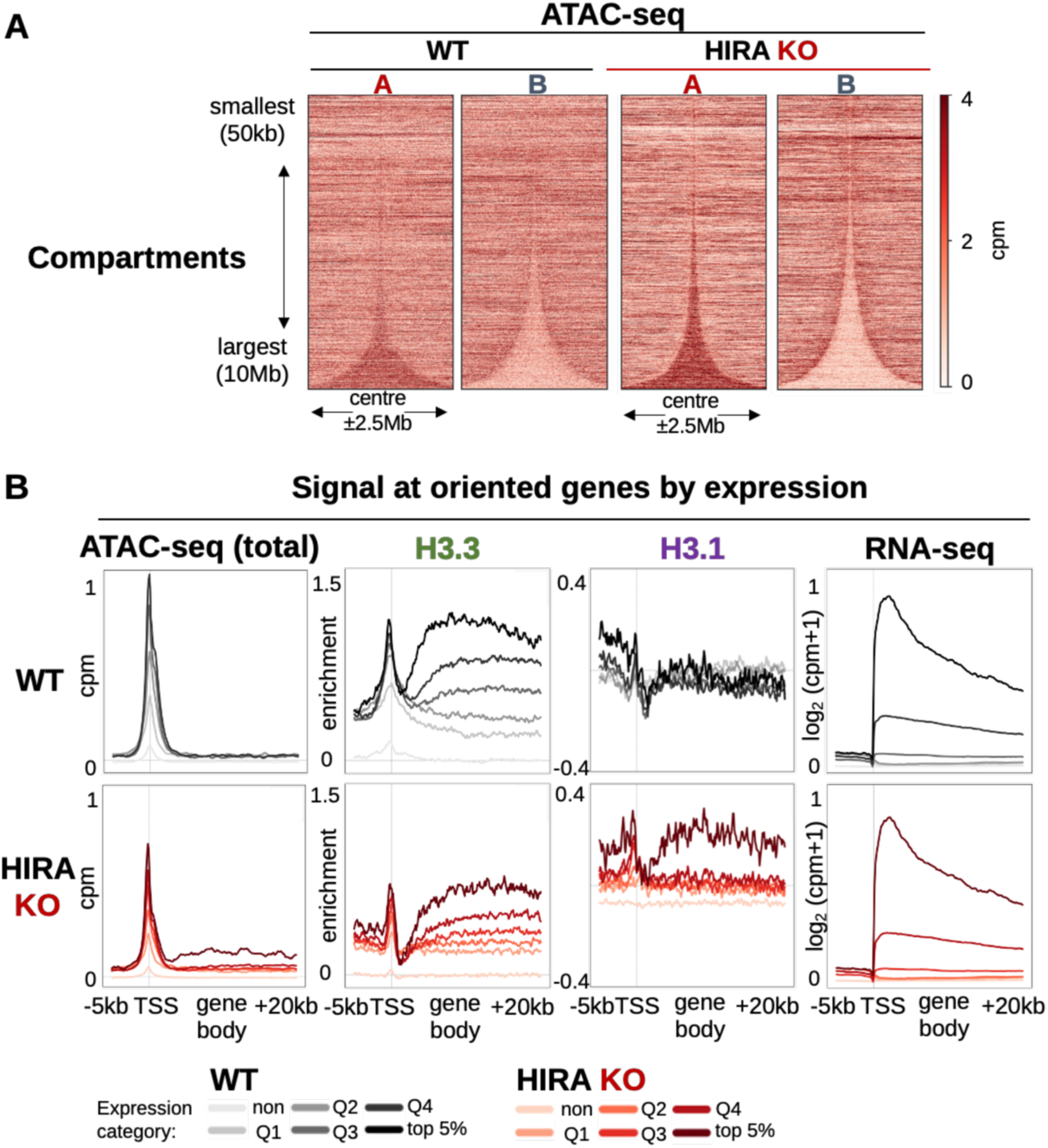
Accessibility increases in compartment A and at active gene bodies in the absence of HIRA. **A.** ATAC-seq at 10kb bins at compartments A and B from WT and HIRA KO HeLa cells, sorted by size and centered at their middle ±2.5Mb. **B.** Total ATAC-seq (no fragment length filtering), H3.3, H3.1 enrichment and RNA-seq binned at 100bp from WT (top) and HIRA KO cells (bottom) at TSSs (−5kb upstream/+20kb downstream). Signals are stratified based on gene expression level from variance-normalised RNA-seq counts obtained from DESeq2. Expression categories: non (undetected, n = 18883), Q1 (1-25% expression, n = 4407), Q2 (26-50% expression, n = 4376), Q3 (51-75% expression, n = 4391), Q4 (76-95% expression, n = 3513) and top 5% expressed genes (n = 879). ATAC-seq is shown as cpm. H3.3 and H3.1 enrichment shown is z-score of log_2_ IP/input. RNA-seq is shown as log_2_(cpm+1). Genes were stratified based on expression level from variance-normalised RNA-seq counts obtained from DESeq2.

### Local contacts and insulation of self-interacting domains demarcated by active genes decrease along with accessibility changes

We further probed the link between HIRA-mediated nucleosome assembly and higher-order chromatin organisation. We stratified the active genes by expression as above and examined local chromatin organisation in their vicinity. Genes are known to insulate contacts between flanking region by possibly serving as extrusion barriers. Two distinct mechanisms have been considered: (i) for genes colocalising with CTCF, CTCF which can halt extrusion and thus establish the borders of domains (Dixon et al., 2012; Hsieh et al., 2020), and (ii) even in the absence of CTCF, active genes can pause extrusion by themselves through the “moving barrier” mechanism (Banigan et al., 2023). In WT cells, we observe the strongest insulation at highly transcribed genes (Figure 5). In the absence of HIRA, this insulation of gene-flanking regions within a range of 0.1-1Mb distance from active genes strongly decreased (Figure 5, Supplementary Figure 5A). These data are in line with the global reduction in compartment A interactions (Figure 2A-C) and its increased accessibility (Figure 4A). The fact that reduced insulation by active genes directly correlated to their level of expression put forward a transcription-mediated interference as a key mechanism (Figure 5). Given this observation, we wondered whether the function of other known insulators was also affected in the absence of HIRA. In this context, CTCF sites were particularly attractive to consider. Indeed, they are enriched at the borders of self-interacting domains (Dixon et al., 2012) and involved in their insulation (Nora et al., 2017; Wutz et al., 2017). Furthermore, CTCF sites are known to be enriched in H3.3 (Jin et al., 2009; Lacoste et al., 2014) in a HIRA-dependent manner (Supplementary Figure 2A). Remarkably, we found reduced insulation only at CTCF sites associated with an active TSSs (Supplementary Figure 5B). We thus conclude that the reduction of insulation observed in HIRA KO depends on a transcription-associated mechanism driven by active genes.

**Figure 5.**
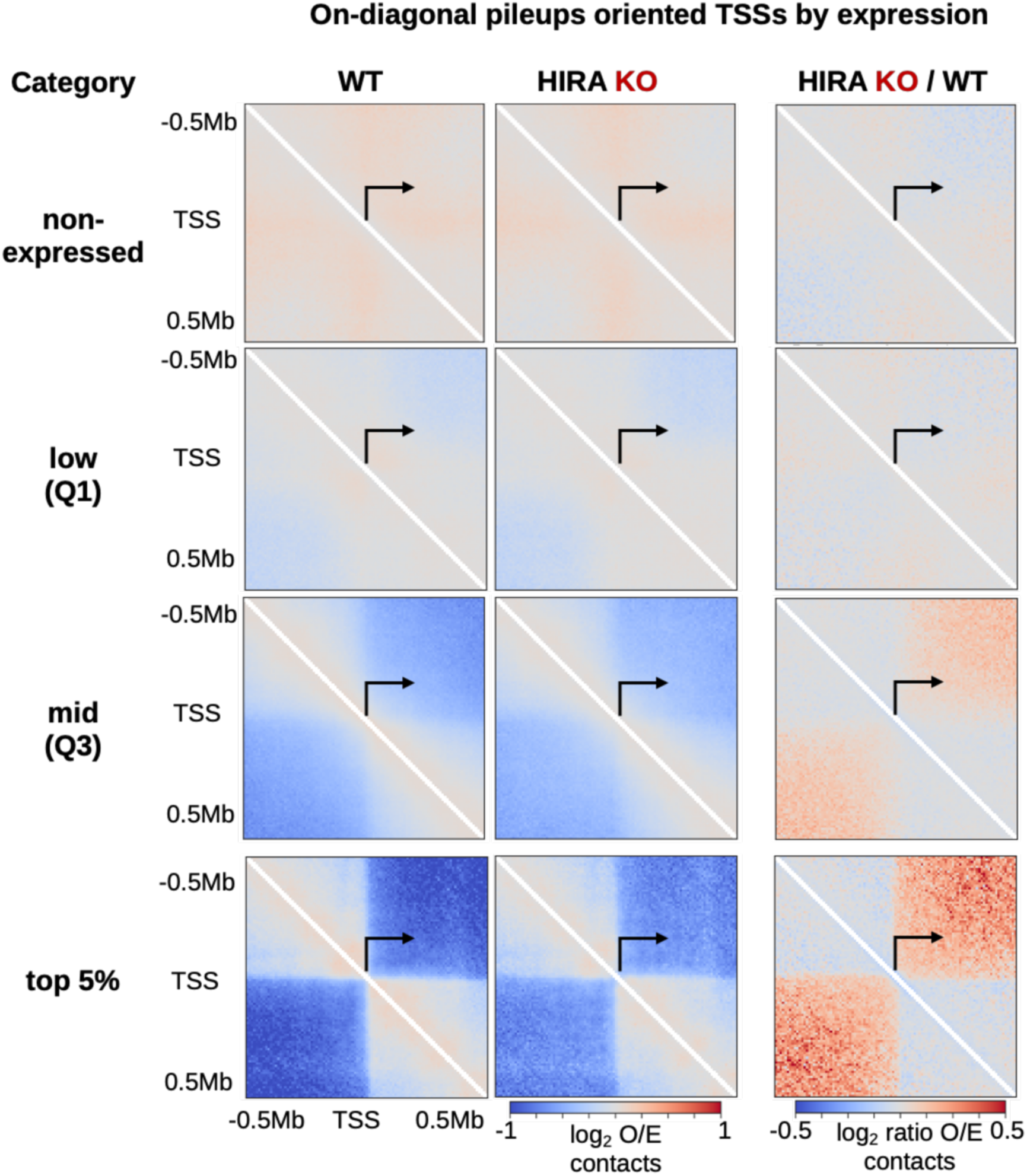
In the absence of HIRA insulation by active genes decrease, scaling with expression level On-diagonal pileups at oriented TSS (±0.5Mb) from WT and HIRA KO (left) and their differential (right, log_2_ HIRA KO/WT). Notice insulation created by highly transcribed genes, and reduction of such insulation in HIRA KO at top genes. All data are at 10kb-binned observed/expected (O/E) Hi-C. Genes were stratified based on expression from variance-normalised RNA-seq counts obtained from DESeq2. Expression categories: non-expressed (undetected, n = 18883), low/Q1 (1-25% expression, n = 4407), mid/Q3 (51-75% expression, n = 4391) and top 5% expressed genes (n = 879), as in Figure 4. Arrows denote TSS position and orientation.

## Discussion

In this study, we explored how HIRA-dependent H3.3 deposition impacts higher-order chromatin folding. Using the knock-out (KO) of the histone chaperone HIRA combined with integrating Hi-C, ChIP-seq and ATAC-seq data, we revealed that impaired H3.3 deposition in active chromatin via this distinct pathway impacts chromatin folding across scales. In our model, targeted deposition of H3.3 by HIRA ensures a link between the nucleosome- and the compartment-level organisation (Figure 6). We propose that HIRA contributes to the local folding and compartment organisation of active chromatin by maintaining H3.3 nucleosomes at expressed genes through redeposition of nucleosomes lost upon transcription-mediated eviction. We discussed here three major aspects: (i) the importance of HIRA-mediated H3.3 incorporation for compartment A interactions, (ii) its link to the local organisation of active genes, and (iii) how these may relate to the importance of HIRA for cell fate transitions in different physiological contexts.

**Figure 6.**
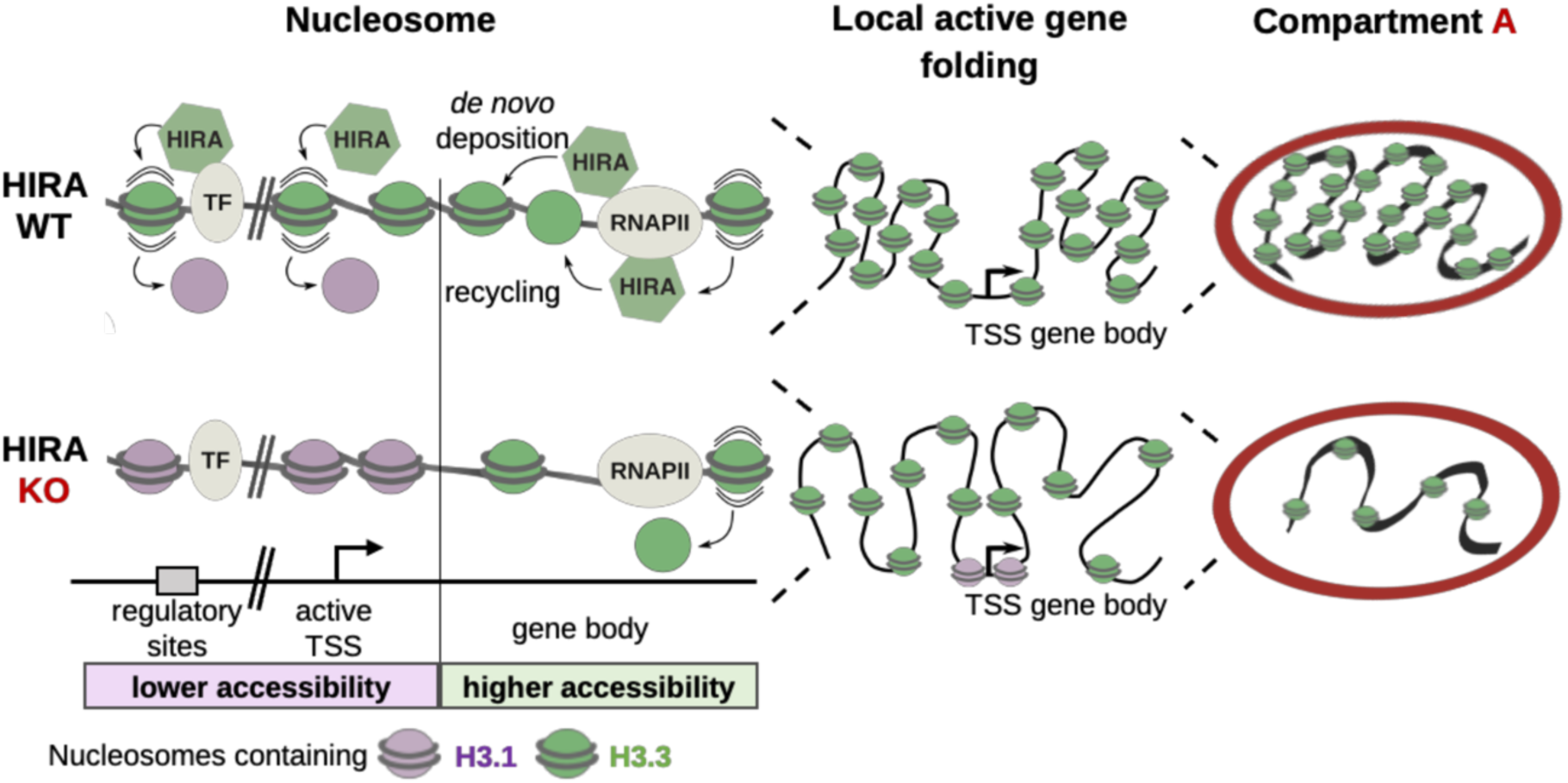
Model for the contribution of HIRA-mediated nucleosome assembly at active genes to their local 3D organisation and compartment A interactions Top: In WT cells, HIRA deposits H3.3 at regulatory elements and active TSSs, promoting their accessibility, and at active genes, maintaining their nucleosome density. Locally, this supports insulation of self-interacting domains by active genes. On the compartment scale, it results in enrichment of H3.3 in compartment A. Bottom: In the absence of HIRA, regulatory sites and active TSSs show lower H3.3 and higher H3.1 enrichment along with reduced accessibility, indicating nucleosome turnover may be reduced. In contrast, decreased H3.3 deposition at active gene bodies is accompanied by increased accessibility. Locally, this is associated with reduced local contacts and insulation of self-interacting domains by active genes. In compartment A, H3.3 enrichment decreases while accessibility increases. The absence of HIRA leads to minor compartment switching and reduces compartment A interactions genome-wide without redistribution of H3 PTMs.

### HIRA-mediated H3.3 nucleosome assembly promotes compartment A contacts independently of PTMs

In the absence of HIRA, H3.3 enrichment decreases in compartment A, without redistribution of H3 PTM enrichment. This was unexpected given the local reduction of H3K4me1 and H3K27ac at corresponding peaks in HIRA KO cells. Indeed, phosphorylation of the unique H3.3S31 residue promotes p300 activity and increase H3K27ac at regulatory elements (Martire et al., 2019; Morozov et al., 2023). In contrast, loss of the H3K4me1-specific methylases MLL3/4 results in reduced H3.3 at H3K4me1 peaks in mouse embryonic stem cells (Goekbuget et al., 2023). This data highlights that although H3.3 and enhancer-associated PTMs may be functionally linked at regulatory regions, changes in their distribution at this genomic scale does not necessarily translate into reorganisation with respect to larger domains. Previous work reported that modifications on both H3.1 and H3.3 from oligonucleosomes show comparable features and rather relate to the chromatin environment (Loyola et al., 2006). Therefore, we conclude that at the scale of compartments, the pre-existing chromatin environment and not the choice of H3 variant dictate H3 PTM distribution.

Compartment A contacts decreased along with the decrease in H3.3 enrichment in HIRA KO cells (Figure 6, right). Several studies have provided support to the idea that histone PTM states may contribute to compartmentalisation (Falk et al., 2019; Hildebrand and Dekker, 2020), although the mechanism remains unclear (Li et al., 2024). However, H3 PTMs did not redistribute at this scale in our system. In contrast, chromatin accessibility in compartment A increased, likely due to the impaired H3.3 deposition over active gene bodies in the absence of HIRA (Figure 6, left). This suggests that interactions of compartment A organisation are influenced not only by the histone marks, as widely accepted, but also by the nucleosome organisation and density. Such reduction of nucleosome density in active regions, largely gene bodies, can lead to weakening of interactions of compartment A. While mammalian compartmentalization is driven largely by affinities between heterochromatin regions (Falk et al., 2019), recent studies (Aljahani et al., 2022; Friman et al., 2023; Goel et al., 2023) have revealed enrichment of interactions between genes and regulatory regions (broadly H3K27ac regions) suggesting that weak yet non-negligible affinities between them should also be considered. We propose that the decrease in nucleosome density over gene bodies in HIRA KO leads to weaker affinities mediated by interactions between nucleosomes. Overall, our data support a view according to which HIRA-dependent nucleosome assembly in active chromatin contributes to its higher-order organisation, impacting compartment A organisation independently of histone PTMs.

### HIRA controls active gene accessibility and insulation of self-interacting domains that they flank

In WT cells, HIRA promotes H3.3 incorporation in active regions. This results in (i) high turnover at active regulatory sites and TSSs associated with their high accessibility, and (ii) maintenance of nucleosome density over gene bodies, thereby restricting accessibility. Indeed, in the absence of HIRA, both regions lose H3.3 enrichment. However, accessibility decreases at regulatory elements while increasing in active gene bodies (Figure 6, left). The lower accessibility at regulatory elements without HIRA is consistent with previous reports (Martire et al., 2019; Morozov et al., 2023; Tafessu et al., 2023). Here, we show it is linked to an increase of H3.1, suggesting that H3.3 incorporation may create a dynamic environment, promoting accessibility of TSS and regulatory sites. However, the increase of accessibility along active gene bodies was unanticipated. We propose that in the absence of HIRA, H3.1 cannot compensate for transcription-coupled H3.3 eviction over gene bodies. This is in line with a dominant role for the H3.3 incorporation by HIRA during transcription that dramatically affects nucleosome density over gene bodies. The resulting increased accessibility over gene bodies suggests a mechanism of interference relying on a HIRA-dependent histone deposition. These changes in nucleosome density also have an unanticipated impact at a higher scale in chromatin organization.

Indeed, the loss of HIRA results in reduced insulation of regions flanking active genes, while having no effect on insulation by CTCFs or disrupting TADs. Remarkably, the effect correlates perfectly with transcription levels. Together, this suggests that HIRA affects transcription-mediated (CTCF-independent) insulation. Such insulation is believed to result from ability of transcribing RNA Pol II to slow-down cohesin and even push it back, known as “the moving barrier” mechanism seen both in mammals and in bacteria (Banigan et al., 2023; Brandão et al., 2019). How can reduced nucleosome occupancy at gene bodies in HIRA KO interfere with this process and make insulation weaker? An attractive possibility to explain this is an effect of nucleosome density on molecular motors participating in this process: RNA Pol II, cohesin, or both. On the one hand, nucleosomes present barriers to RNA Pol II (Lorch et al., 1987) and hinder its passage. On the other hand, nucleosomal organisation creates a fibre that cohesion extrudes into a loop. Thus, reduced nucleosome density in the absence of HIRA could affect speeds of both cohesin-driven loop extrusion and transcribing polymerases acting as moving barriers (Banigan et al., 2023). Indeed, lower density of nucleosomes would make the same stretch of DNA form a longer DNA-nucleosome fibre. Since an extended fibre would increase the demand for cohesin to extrude the same length of DNA (Maji et al., 2020), the HIRA mediated effect that we observed could build on such properties. Effectively, slower extrusion would then result in weaker insulation. Importantly, HIRA has been involved in both *de novo* H3.3 deposition (Ray-Gallet et al., 2011) and recycling of old H3.3 (Torné et al., 2020) in the context of transcription. Therefore, these data support a significant role for HIRA in maintaining nucleosome occupancy in transcribed regions impacting local interactions and insulation in the 0.1-1Mb range.

In conclusion, our work highlights the importance of H3.3 nucleosome maintenance by HIRA in impacting accessibility and higher-order organisation of active chromatin. Thus, in addition to the previous role of HIRA in promoting accessibility at regulatory peaks (Martire et al., 2019; Morozov et al., 2023; Tafessu et al., 2023), here we demonstrate a key role for HIRA to limit accessibility in active gene bodies. These findings help explain how HIRA promotes insulation by active genes in a manner linked to their accessibility. Given the importance of HIRA for induction of new transcriptional programmes (Banaszynski et al., 2013; Fang et al., 2018; Gomes et al., 2019), or changes in cell states with major subnuclear compartment changes as observed in senescent cells (Kennedy et al., 2010; Lee and Zhang, 2016; Zhang et al., 2005), future work should address which aspects of HIRA function is key in these contexts. Interestingly, in prostate cancer where HIRA expression correlates with worse prognosis (Morozov et al., 2023), recent reports showed compartment A accessibility associated with higher rate of structural variation (Zhao et al., 2024). Thus, our findings reveal a new aspect of HIRA function which may contribute to cancer progression. Finally, we demonstrate that HIRA contributes to compartment A organisation independently of histone marks, disentangling the correlation between H3.3, histone PTMs and A/B compartments. We anticipate that this view can stimulate new avenues to explore the role of histone variant dynamics at distinct sites or developmental stages on higher-order chromatin organisation.

## Materials and Methods

### Cell culture

We used HeLa cells stably expressing H3.1-SNAP-HA or H3.3-SNAP-HA which were either wild-type (WT, CRISPR/Cas9 GFP KO) or knock-out for HIRA (KO, CRISPR/Cas9 HIRA KO), as described in (Ray-Gallet et al., 2018). We cultured and synchronised cells at the G1/S transition by double thymidine block as described in (Forest et al., 2024; Gatto et al., 2022). All cell lines were tested negative for mycoplasma.

### Hi-C

We performed Hi-C using the Arima Hi-C+ kit following the manufacturer’s instructions. Briefly, 5-10 million asynchronous HeLa cells per condition were fixed in 4% formaldehyde for 10min before quenching the reaction with Stop Solution 1. Fixed cells were washed in PBS and snap-frozen in liquid nitrogen at 1 million cell aliquots. We performed cell and nuclear lysis, restriction enzyme digestion, end repair, biotinylation, ligation and decrosslinking as per manufacturer’s instructions and proximally ligated DNA was isolated using AMPure XP beads. Arima QC1 was performed and was successful for all samples. We sonicated DNA using Covaris E220 Evolution (100μL sample, 7°C, peak incident power: 105W, duty factor: 5%, cycles/burst: 200 for 100s), and fragmented DNA was size selected using AMPure XP beads. We verified that the mean size of selected DNA was 400bp using Tapestation. After biotin enrichment, we performed library preparation using the KAPA HyperPrep kit, following the modified protocol described in the Arima Hi-C+ kit instructions. After ligation of Illumina TruSeq sequencing adaptors, we performed Arima QC2 (library quantification) to determine the number of amplification cycles for the library PCR using the KAPA Library Quantification Sample kit following the manufacturer’s instructions. Amplified libraries were sequenced on Illumina NovaSeq 6000 (PE100) at the NGS (Next-Generation Sequencing) platform at Institut Curie.

### Histone PTM ChIP-seq

We performed ChIP-seq of histone post-translational modifications (histone PTM ChIP-seq) using the native nucleosomes isolation procedure described in (Gatto et al., 2022) with small modifications. We used 5 million asynchronous cells per IP and Dynabeads Protein A-conjugated antibodies against histone PTMs for immunoprecipitation. All steps were performed at 4°C and in the presence of Protease inhibitors (Roche) and 1mM TSA in every buffer to prevent protease and HDAC activity, respectively. For each IP reaction, we prepared 50μL of Ab-conjugated beads by blocking for 4 hours in Bead blocking buffer (2.5% BSA in PBS-T, 1mg/mL tRNA), washing once in 0.02% PBS-T and incubating for 15-30min with the antibodies (diluted in 0.2mL 0.02% PBS-T) on a rotating wheel. Ab-conjugated beads were then washed twice with 0.02% PBS-T and resuspended in 0.2mL 0.02% PBS-T. We pooled native nucleosomes (80μL per IP) from up to 3 samples, diluted in 5x volumes of Incubation buffer (50mM Tris-HCl pH 7.5, 100mM NaCl, 0.5% BSA) and pre-cleared by incubating with Dynabeads Protein A (15μL per input pool) for 30 min on a rotating wheel. We kept 20μL (1%) pre-cleared chromatin as input sample. We incubated the remaining (460μL/IP) with the Ab-conjugated beads (washed once in Incubation buffer) overnight on a rotating wheel and purified DNA as described for SNAP-seq in Gatto et al. (2022). Sequencing libraries were prepared at the NGS (Next-Generation Sequencing) platform at Institut Curie with the Illumina TruSeq ChIP kit and sequenced on Illumina NovaSeq 6000 (PE100).

### ATAC-seq

To assay accessibility, we performed ATAC-seq from 50000 G1/S-synchronised cells per condition using the Active Motif ATAC-seq kit (53150) following the manufacturer’s instructions. We verified DNA profiles before sequencing on Illumina NovaSeq 6000 (PE100) by the NGS platform at Institut Curie.

### Total RNA-seq

We obtained total RNA from 1 million G1/S-synchronised cells per condition. We collected cells by trypsinisation and extracted RNA with the RNeasy Plus Mini Kit (QIAGEN) including DNase treatment (RNase-free DNase QIAGEN, 79254) using manufacturer’s instructions. We quantified RNA using Nanodrop and checked the quality by Tapestation. We used 10ng of total RNA for library preparation using TruSeq Stranded Total RNA kit and sequenced libraries on Illumina NovaSeq 6000 (PE100) at the NGS platform at Institut Curie.

### ChIP-seq, ATAC-seq and RNA-seq data processing

ChIP-seq and ATAC-seq data was processed from raw reads in FASTQ format as described in (Gatto et al., 2022). Briefly, for each sample, we mapped reads to the soft-masked human reference genome (GRCh38) downloaded from Ensembl (release 109) using bowtie2 v2.3.4.2 (Langmead and Salzberg, 2012) with --very-sensitive parameters. RNA-seq data was aligned with hisat2 v2.1.0 (Kim et al., 2019), run in paired-end mode with default parameters. We used SAMtools v1.9 (Danecek et al., 2021) to sort, flag duplicates and index bam files for all samples. We used samtools view (-f 2 -F 3840 parameters to keep reads mapped in pairs and exclude QC fails, non-primary alignments and duplicates) to compute coverage over the genome as a BED file (chromosome, start, end, MAPQ). Quality control of ATAC-seq data was additionally performed on bam files by ATACseqQC v1.28.0 (Ou et al., 2018). To obtain non-nucleosomal reads for the ATAC-seq data, we filtered fragments of <125bp length from the BAM files using samtools. Then, we used BEDtools v2.27.1 (Quinlan and Hall, 2010) to calculate number of fragments in consecutive 100bp or 1kb bins. To obtain gene-level counts from RNA-seq data, we used featureCounts (Subread v1.6.3) (Liao et al., 2014) in paired-end mode, filtering for MAPQ>2. We aligned reads on the sequences of the ERCC92 RNA spike-in control from Life Technologies.

Analysis of the data was carried out by custom Python scripts using pandas v1.5.3 (McKinney, 2010), NumPy v1.23.5 (Harris et al., 2020) and scipy v1.11.2 (Virtanen et al., 2020). Visualization was performed using matplotlib v3.6.2 (Hunter, 2007) and seaborn v0.12.2 (Waskom, 2021). For each sample, counts were read at 100bp or at 1kb resolution and aggregated to 10kb bins. Binned data (at 100bp and 10kb) was normalized to the total number of mapped counts to generate cpm (counts per million). ChIP-seq data was then normalized to matched input sample by computing a log_2_ ratio (IP/input). To enable cross-sample comparison, ChIP-seq signal was then scaled and centered by computing a z-score per chromosome. RNA-seq data was plotted as log_2_(cpm+1). Gene annotation was obtained from Ensembl (v99), CTCF site locations in HeLa were obtained from ENCODE (ENCFF816CUG). CTCF sites were classified as TSS-far (>50kb from a H3K4me3 peak, not in gene bodies, in compartment A) and TSS-near (<1kb from an annotates TSS, overlapping a H3K4me3 peak, in compartment A), using H3K4me3 peak calls (described below) and compartment call from WT cells.

### Peak calling

Peaks for active PTMs (H3K4me1, H3K4me3 and H3K27ac) were called using MACS2 v2.2.7.1 callpeak (Gaspar, 2018; Zhang et al., 2008) from bam files using matched inputs as background (-c) for each condition and bandwidth was set to 400 (--bw). Common peaks between the two cell lines for each mark and condition were obtained by overlap using bioframe v0.4.1 (Open2C et al., 2022) and used for downstream analysis. ATAC-seq peaks were called from each replicate and condition using HMMRATAC (Tarbell and Liu, 2019) with default parameters except --window 2500000. Common peaks between the two replicates and cell lines for each condition were obtained by bioframe.overlap used for downstream analysis.

### RNA-seq analysis

Raw RNA-seq counts were filtered for low detection (retained genes with at least 10 counts over all replicates) and normalized by variance-stabilising transformation (vst) in DESeq2 v1.44.0 (Love et al., 2014). These were used to classify genes by expression level into non-expressed, Q1 (1-25%), Q2 (26-50%), Q3 (51-75%), Q4 (76-95%) and top 5% expressed genes. We performed differential expression between WT and HIRA KO using DESeq2 with lfcThreshold=1 and alpha=0.05 for each cell line and considered the common differentially expressed genes in the two cell lines.

### Hi-C data processing

We used HiC-Pro v3.1.0 (Servant et al., 2015) with default parameters (except MIN_MAPQ=2) to generate raw count Hi-C matrices at 1Mb, 100kb, 50kb, 25kb, 10kb and 5kb resolution from raw FASTQ files. First, we used MultiQC v1.11 (Ewels et al., 2016) to perform quality control and extract the number of short-range (<=20kb), long-range (>20kb) *cis* and *trans* interactions for each sample. HiCExplorer v3.7.2 (Ramírez et al., 2018; Wolff et al., 2020, 2018) was then used to convert matrices to cool format and generate a single mcool file per sample, containing all resolutions listed above.

Matrices were visualized interactively with HiGlass v0.8.0 (Kerpedjiev et al., 2018) at 1Mb and 100kb resolutions to be manually inspected for large inter- and intra-chromosomal aberrations (translocations, inversions, duplications, etc.), and the subsequently generated list of regions was merged with the set of blacklisted regions of the human genome (ensembl). This custom set of blacklisted regions was used to mask matrices prior to normalization by iterative correction (ICE) (Imakaev et al., 2012) with a single iteration using cooler v0.9.3 (Abdennur and Mirny, 2020). Matrix similarity was computed per chromosome at 1Mb, 100kb, 50kb and 10kb resolution with HiCRep v0.2.6 (Lin et al., 2021; Yang et al., 2017) before and after masking without substantial changes. Expected interactions per chromosome arm (coordinates downloaded with bioframe) were calculated using cooltools v0.6.1 (Open 2C et al., 2022). P(s) curves, representing decay of interaction frequency with increasing genomic distance, were calculated per chromosome arm at 10kb resolution, aggregated and smoothed. Analysis was performed in both cell lines, and it gave rise to similar results. Coordinates and data from H3.1-SNAP cells were used for representative images where experiments were performed in both cell lines unless mentioned otherwise.

### Hi-C data analysis

Compartment analysis of Hi-C matrices was performed by eigenvector (EV) decomposition (Lieberman-Aiden et al., 2009) at 50kb resolution with cooltools using GC content track to orient the sign of the first eigenvector (EV1). A/B compartment domains were defined for each sample as contiguous segments of the genome with the same EV1 sign. Compartment switching was determined on a per bin basis, where a compartment-switching bin undergoes the same EV1 sign change in H3.1-SNAP and H3.3-SNAP cell lines from HIRA WT to KO (WT-to-KO). As control, we compared compartment switching between H3.1-SNAP and H3.3-SNAP cells in the same way. Proportion of the genome switching compartment was calculated as the % compartment-switching 50kb bins out of all non-masked 50kb bins. Differential maps to visualize changes in compartment interactions were plotted as log_2_ ratio of ICE-normalized contacts from HIRA KO/WT at 1Mb resolution. To assess A/B compartment interactions genome-wide, we performed saddle-plot analysis using cooltools. For this purpose, we split 50kb-binned EV1 signal into percentiles, re-ordered and averaged O/E-normalised Hi-C maps per chromosome arm from lowest (most strongly B) to highest (most strongly A) EV1 percentile and plotted O/E interaction frequency at 50×50 bins. To generate differential saddle-plots, we calculated the log_2_ ratio of HIRA KO/WT. To quantify the difference in compartment interactions, we split the differential saddle data in six groups, corresponding to interactions between A-A, A-I, A-B, B-B, B-I and I-I, where A, I and B corresponded to the top, middle and bottom tercile of each axis (shown in Figure 2C). TAD borders were identified based on insulation score (Crane et al., 2015), computed from 10kb binned matrices with a window size of 100kb using cooltools. Overlap between border bins (+/-20kb) was determined using bioframe between the two cell lines and between the two conditions. To examine local interactions, on-diagonal pileups were performed from 10kb-binned O/E-normalised matrices oriented in the TSS/CTCF site direction using cooltools.

### Quantifications and statistical analysis

Statistical analysis was performed in pyhton using scipy. p-values were calculated by two-tailed Mann-Whitney U test. Multiple testing correction was performed by controlling the false discovery rate (FDR) using the Benjamini-Hochberg method. Differences with adjusted p-value <0.05 were considered statistically significant.

---

**Supplementary Figure 1.**
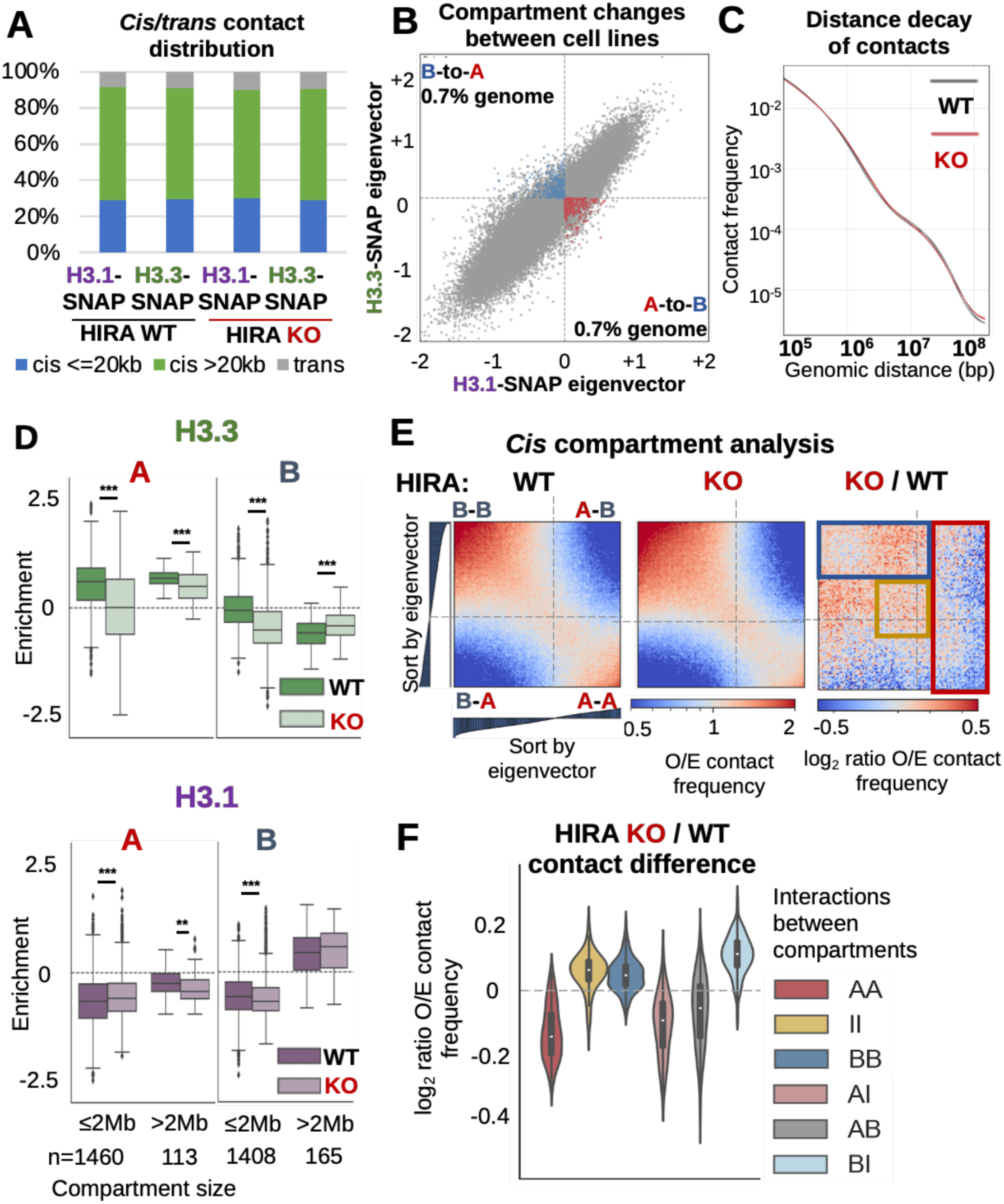
Absence of HIRA leads to re-distribution of H3.3 from A to large B compartments A. Proportion of short-range (<=20kb), long-range (>20kb) *cis* and *trans* contacts from Hi-C maps of HIRA WT or KO H3.1- and H3.3-SNAP HeLa cells. B. EV1 (1^st^ eigenvector, indicating compartment) of 50kb-binned Hi-C matrices from H3.1-vs H3.3-SNAP HIRA WT cells. Bins which change from A-to-B (lower right quadrant) or B-to-A (upper left quadrant) in the same direction in both conditions are coloured red and blue, respectively. C. Decay of contact frequency with genomic distance (P(s) curves) from HIRA WT (grey) and KO (red) maps, binned at 10kb after masking of blacklisted regions and ICE normalization. D. H3.3 and H3.1 enrichment from WT and HIRA KO cells quantified in A/B compartments per indicated domain sizes. Two-tailed Mann-Whitney U test adjusted for multiple testing by FDR (5% cut-off) was used to determine significance of differences between WT and KO. E. Saddle plots from HIRA WT and KO (O/E contacts) and differential (log_2_ ratio of O/E contacts) of HIRA KO/WT at 50kb resolution based on EV1 percentiles from H3.3-SNAP cells. Red quadrangle denotes decreased contacts of compartment A genome-wide, blue quadrangle denotes compartment B interactions with itself and intermediate (I) compartment, and yellow square denotes I-I contacts. F. Difference (log_2_ HIRA KO/WT ratio) of O/E contact frequency between the sets of A/B/I compartments defined in Fig. 2B (scheme) based on differential saddle plots from H3.3-SNAP cells above (Supp. Fig. 1E). H3.3 and H3.1 enrichment is shown as z-score of log_2_ IP/input at 10kb bins. Significance (2-tailed Mann-Whitney U test): ns, p>0.05; *, p<=0.05; **, p<=0.01; ***, p<=0.001

**Supplementary Figure 2.**
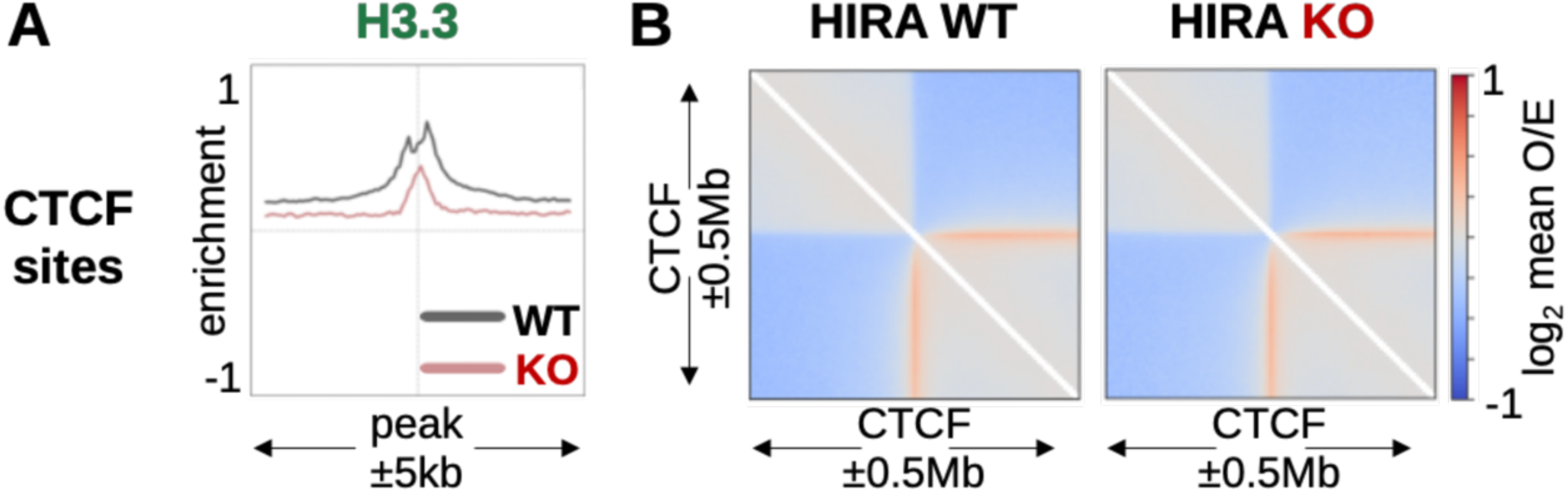
HIRA KO disrupts H3 variant pattern at CTCF sites without impairing their insulation **A.** H3.3 enrichment from WT and HIRA KO cells plotted at oriented CTCF sites (n = 36508) centered at the peak summit ±5kb at 100bp bins. **B.** On-diagonal pileups at oriented CTCF sites ±0.5Mb from 10kb-binned Hi-C of WT and HIRA KO cells. H3.3 enrichment is plotted as z-score of log_2_ IP/input.

**Supplementary Figure 3.**
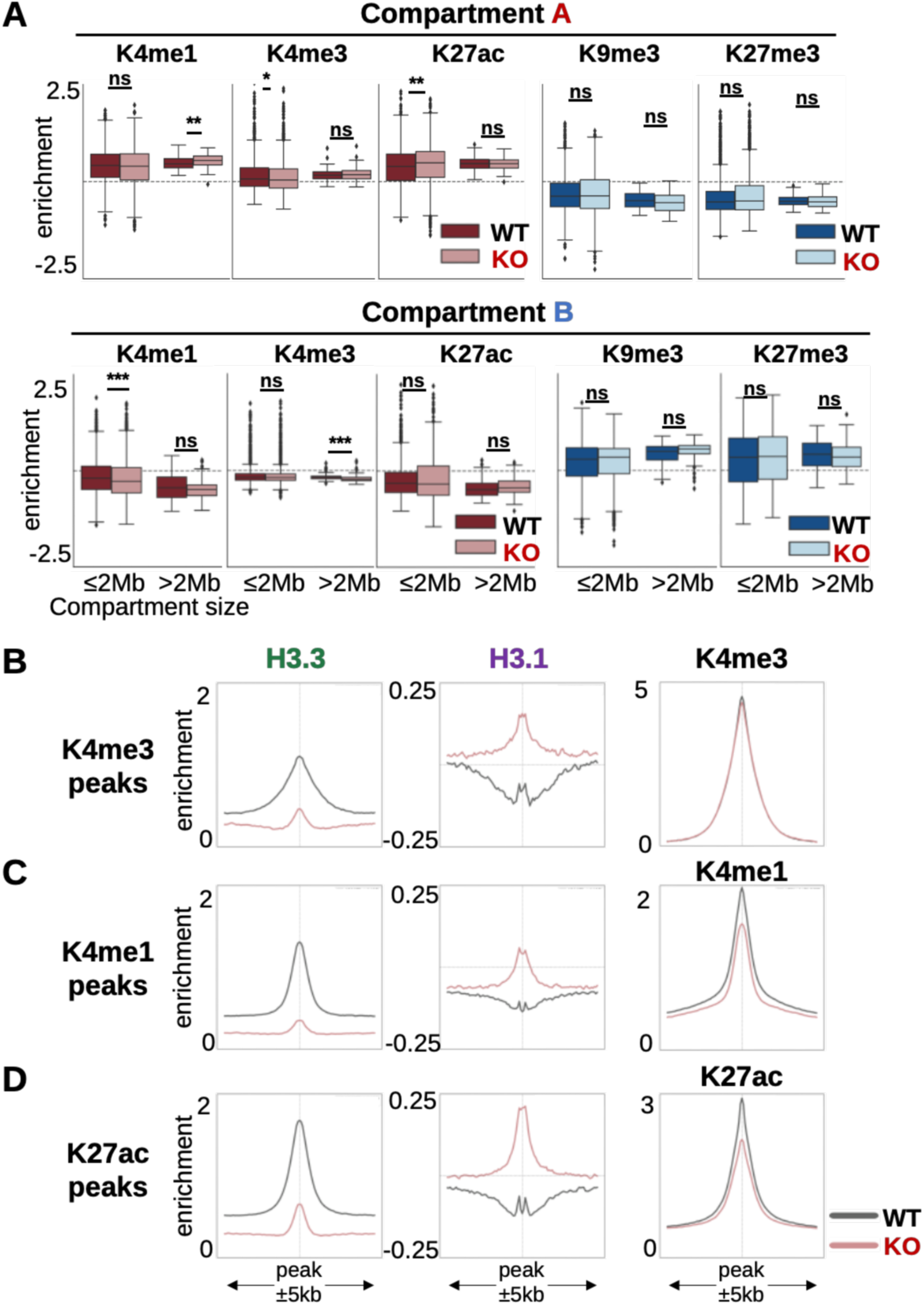
H3.3 redistribution in the absence of HIRA is not accompanied by corresponding PTM changes at the compartment scale **A.** Active (H3K4me1/3, H3K27ac) and repressive (H3K9/27me3) histone PTM enrichment at 10kb bins from HIRA WT and KO cells quantified in compartment A (top) and B (bottom) per indicated domain sizes. Two-tailed Mann-Whitney U test adjusted for multiple testing by FDR (5% cut-off) was used to determine significance of differences between WT and KO. Significance was noted as: * (p<=0.05), ** (p<=0.01), *** (p<=0.001). **B.** Mean H3.3, H3.1 and H3K4me3 enrichment at 100bp bins from WT and HIRA KO at H3K4me3 (n = 26043, promoter-associated) peaks from WT HeLa cells ±5kb. **C.** Mean H3.3, H3.1 and H3K4me3 enrichment at 100bp bins from WT and HIRA KO at H3K4me1 (n = 130608, enhancer-associated) peaks from WT HeLa cells ±5kb. **D.** Mean H3.3, H3.1 and H3K4me3 enrichment at 100bp bins from WT and HIRA KO at H3K27ac (n = 73023, enhancer-associated) peaks from WT HeLa cells ±5kb ATAC-signal is shown as cpm, H3.3, H3.1 and PTM enrichment is shown as z-score of log_2_ IP/input.

**Supplementary Figure 4.**
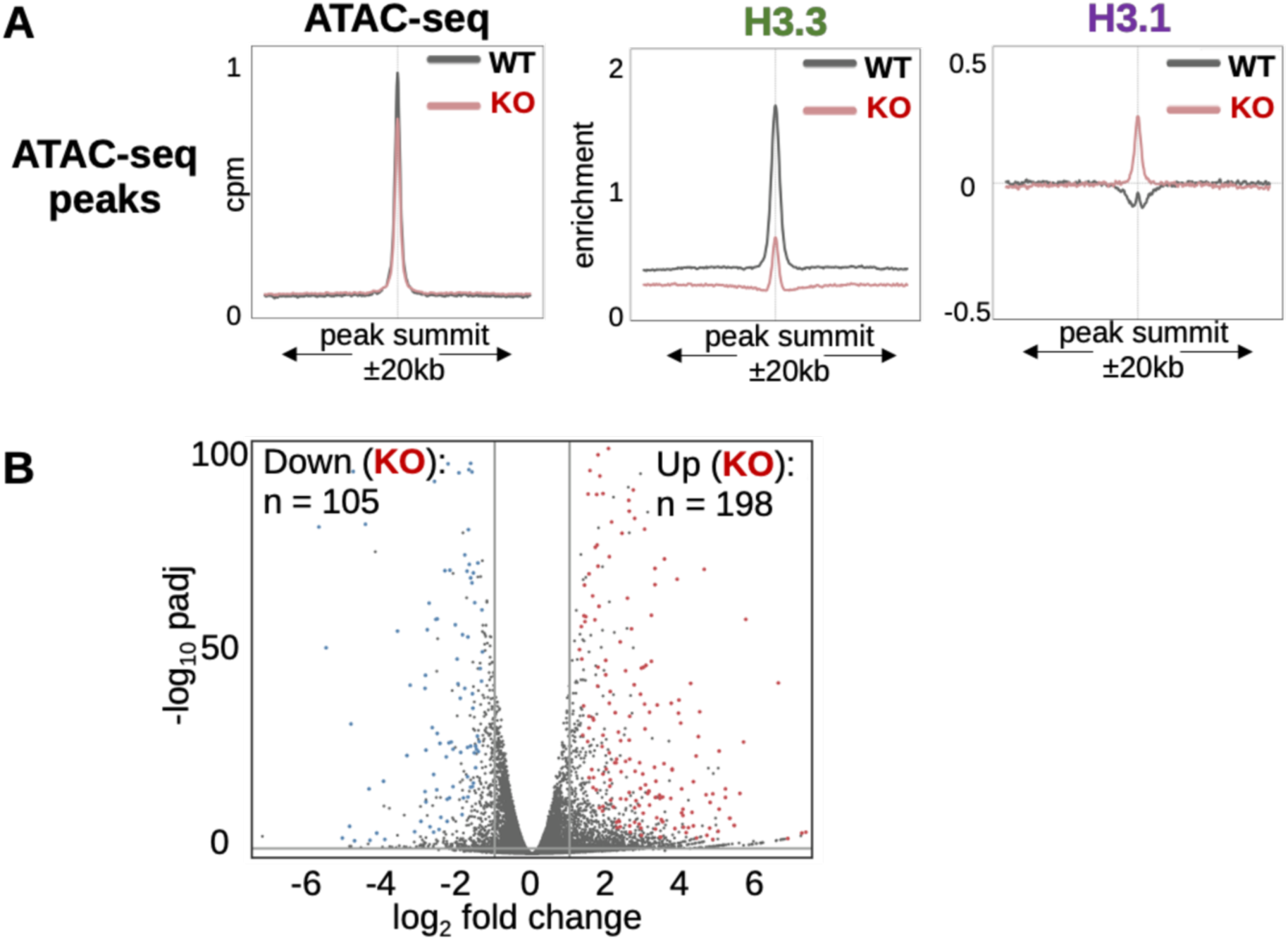
In the absence of HIRA, accessibility changes without differential expression A. ATAC-seq signal, H3.3 and H3.1 enrichment binned at 100bp from WT and HIRA KO at ATAC-seq peaks from WT HeLa cells (n = 45630) ±20kb. B. Volcano plot showing log_2_ fold change (HIRA KO/WT) compared to significance (-log_10_ padj) of expressed genes. Up- and down-regulated genes common for the two cell lines are shown in blue, and red, respectively, and their numbers are noted. Significance was determined using DESeq2 with padj < 0.05 and log_2_ fold change >= 1. ATAC-seq is shown as cpm. H3.3 and H3.1 enrichment are shown as z-score of log_2_ IP/input. RNA-seq is shown as log_2_(cpm + 1).

**Supplementary Figure 5.**
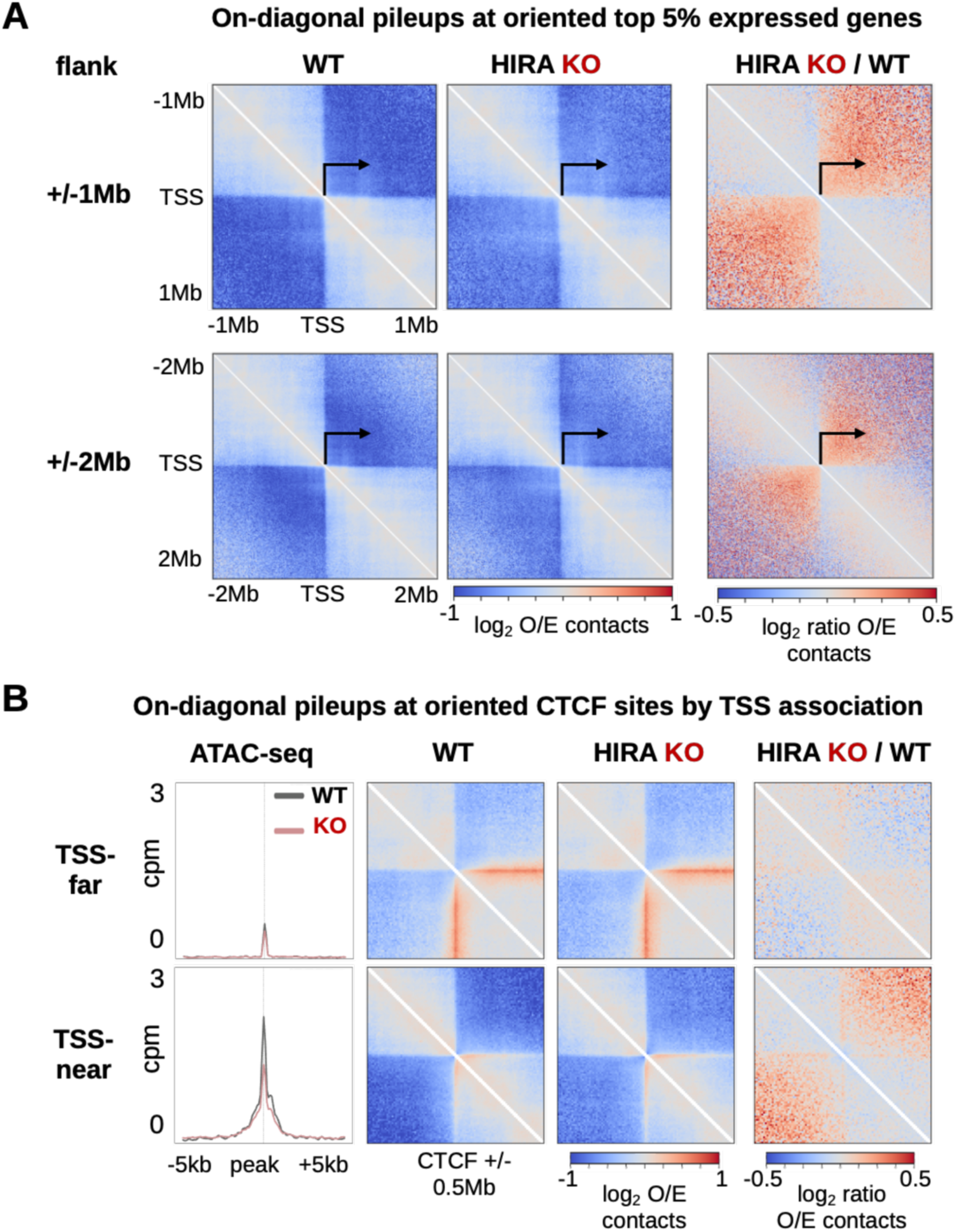
In the absence of HIRA, local contacts and insulation are reduced only in association with gene transcription **A.** On-diagonal pileups at oriented TSSs from to 5% expressed (n = 879) genes, centered at the TSS at different flanking distances (±1Mb, top or ±2Mb, bottom) from 10kb-binned observed/expected (O/E) Hi-C of WT and HIRA KO cells and differential (log_2_ HIRA KO/WT). **B.** ATAC-seq signal and on-diagonal pileups from WT and HIRA KO cells at CTCF sites far from (n = 997) or associated with (n = 721) active TSSs in compartment A. ATAC-seq signal is plotted at CTCF sites ±5kb while pileups are centered on CTCF sites ±0.5Mb. ATAC-seq is binned at 100bp and shown as cpm in grey for WT and in red for HIRA KO. Hi-C matrices were observed/expected (O/E) normalised at 10kb resolution. Differential pileups are shown as log_2_ HIRA KO/WT. Non-expressed genes were selected based on gene expression level from variance-normalised RNA-seq counts from DESeq2.

**Supplementary Table 1.** Detection of TAD borders between cell line and condition.

| Detected in both cell lines | Detected in both conditions | Number of TAD borders |  |  |  | Percentage of TAD borders |  |  |  |
| --- | --- | --- | --- | --- | --- | --- | --- | --- | --- |
|  |  | H3.1 WT | H3.3 WT | H3.1 KO | H3.3 KO | H3.1 WT | H3.3 WT | H3.1 KO | H3.3 KO |
| yes | yes | 5413 | 5381 | 5385 | 5439 | 84,78 | 80,53 | 81,46 | 84,01 |
| yes | no | 411 | 443 | 489 | 435 | 6,44 | 6,63 | 7,4 | 6,72 |
| no | yes | 276 | 368 | 302 | 309 | 4,32 | 5,51 | 4,57 | 4,77 |
| no | no | 285 | 490 | 435 | 291 | 4,46 | 7,33 | 6,58 | 4,49 |
| total |  | 6385 | 6682 | 6611 | 6474 |  |  |  |  |

**Supplementary Table 2.** Overlap of active PTM peaks common between cell lines between WT and HIRA KO.

| Sample | Number of PTM peaks |  |  | Percentage of PTM peaks |  |
| --- | --- | --- | --- | --- | --- |
|  | all | shared (WT & KO) | unique (WT or KO) | shared (WT & KO) | unique (WT or KO) |
| K4me1_WT_r1 | 130608 | 101708 | 28900 | 77,87 | 22,13 |
| K4me1_WT_r2 | 131307 | 113765 | 17542 | 86,64 | 13,36 |
| K4me1_KO_r1 | 163275 | 131314 | 31961 | 80,43 | 19,57 |
| K4me1_KO_r2 | 141424 | 102066 | 39358 | 72,17 | 27,83 |
| K4me3_WT_r1 | 26043 | 23544 | 2499 | 90,4 | 9,6 |
| K4me3_WT_r2 | 23255 | 22580 | 675 | 97,1 | 2,9 |
| K4me3_KO_r1 | 25255 | 23688 | 1567 | 93,8 | 6,2 |
| K4me3_KO_r2 | 23270 | 20949 | 2321 | 90,03 | 9,97 |
| K27ac_WT_r1 | 73023 | 49198 | 23825 | 67,37 | 32,63 |
| K27ac_WT_r2 | 66746 | 52981 | 13765 | 79,38 | 20,62 |
| K27ac_KO_r1 | 69793 | 57425 | 12368 | 82,28 | 17,72 |
| K27ac_KO_r2 | 64915 | 48084 | 16831 | 74,07 | 25,93 |

**Supplementary Table 3.** Overlap of ATAC-seq peaks common between cell lines and replicates between WT and HIRA KO.

| Sample | Number of ATAC-seq peaks |  |  | Percentage of ATAC-seq peaks |  |
| --- | --- | --- | --- | --- | --- |
|  | all | shared (WT & KO) | unique (WT or KO) | shared (WT & KO) | unique (WT or KO) |
| H31_WT_r1 | 45630 | 41586 | 4044 | 91,14 | 8,86 |
| H31_WT_r2 | 46777 | 44387 | 2390 | 94,89 | 5,11 |
| H31_KO_r1 | 56885 | 42478 | 14407 | 74,67 | 25,33 |
| H31_KO_r2 | 61588 | 47113 | 14475 | 76,5 | 23,5 |
| H33_WT_r1 | 50357 | 44328 | 6029 | 88,03 | 11,97 |
| H33_WT_r2 | 54418 | 48694 | 5724 | 89,48 | 10,52 |
| H33_KO_r1 | 57287 | 49898 | 7389 | 87,1 | 12,9 |
| H33_KO_r2 | 62913 | 53949 | 8964 | 85,75 | 14,25 |

## Materials

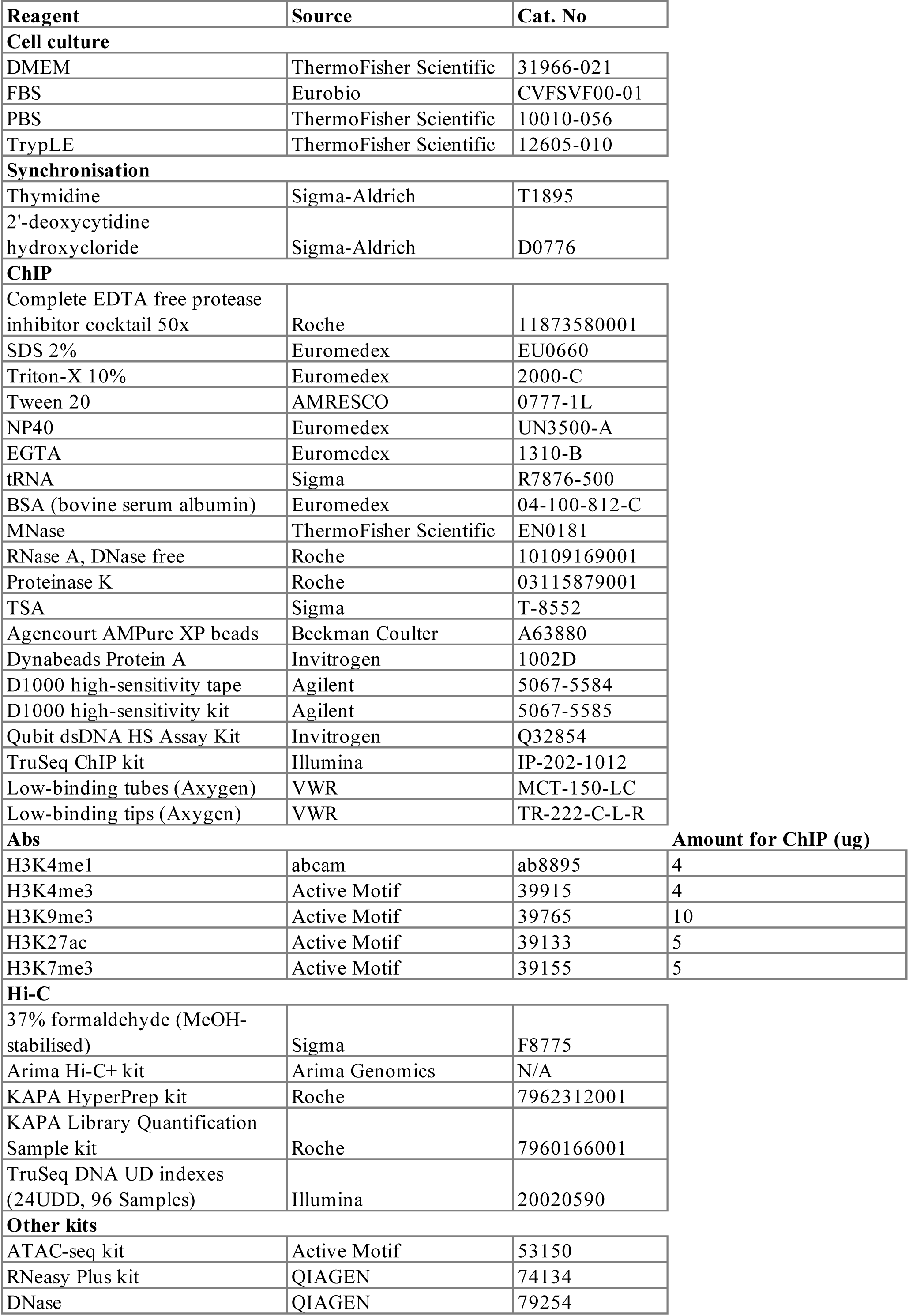
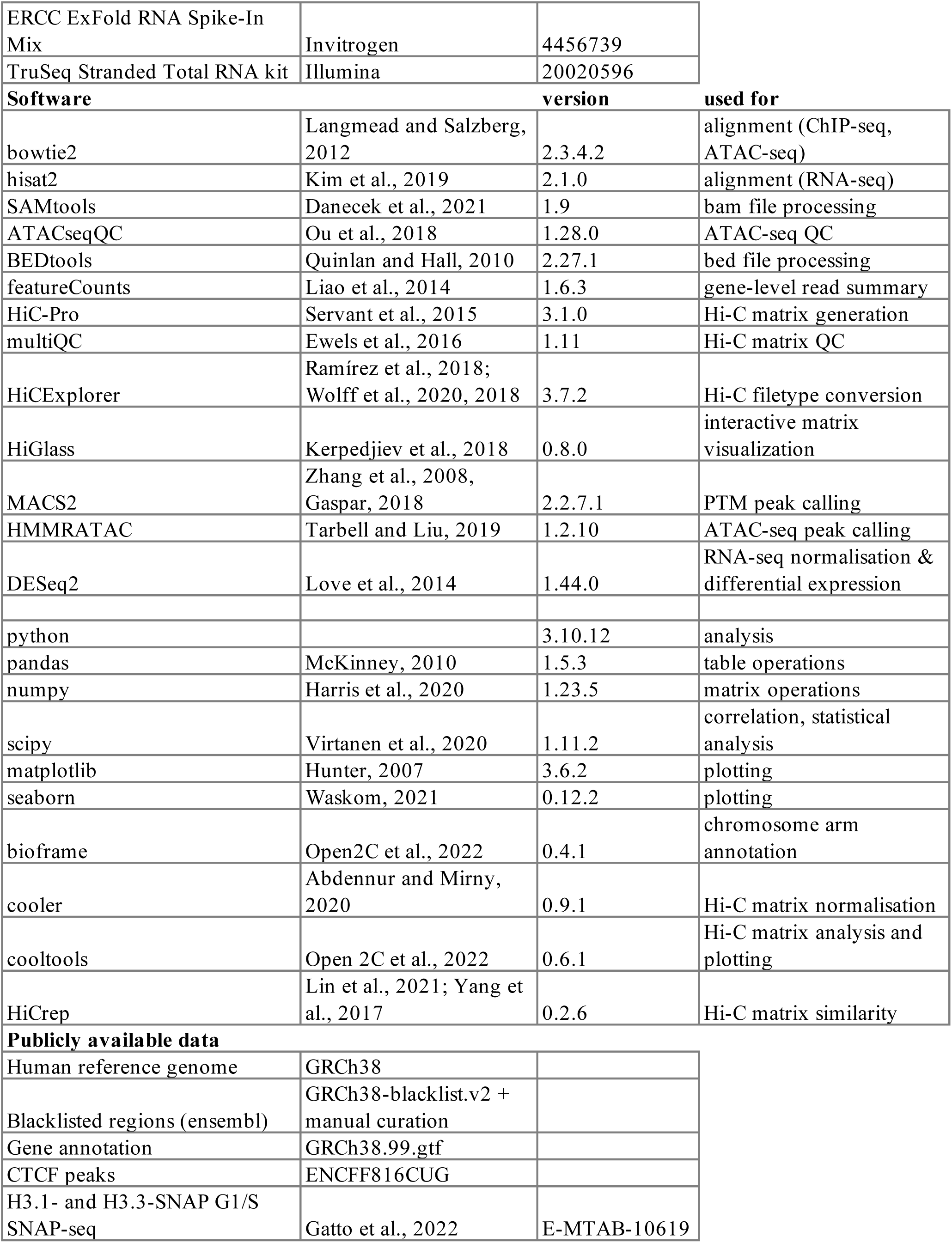

### Author contributions

G. A. and T.K. conceived the overall strategy. T.K. and G.A. wrote the paper. T.K., A.F. and J.P.Q. performed the experiments. T.K. generated most of the figures and analysed data. G.A. supervised the work. M.A.M-R., A.G. and L.M. provided advice for the analysis. Critical reading and discussion of data involved all authors.

## Acknowledgements

We thank the members of UMR3664 and Almouzni team for helpful discussions. We thank Sébastien Lemaire for critical reading of the Methods, Dominique Ray-Gallet for constructive feedback on the model, Héloïse Muller and Nicolas Servant for advice on the Hi-C experiments and analysis. We acknowledge the ICGex NGS platform of the Institut Curie. This work was supported by the European Research Council (ERC-2015-ADG-694694 ‘‘ChromADICT’’), the Ligue Nationale contre le Cancer (Equipe labellisée Ligue), France and Agence Nationale de la Recherche, France (ANR-11-LABX-0044_DEEP, ANR-10-IDEX-0001-02 PSL, and ANR21-CE-11-0027 ‘‘CAFinDs’’). T.K. was supported by individual funding from H2020 MSCA-ITN – ChromDesign (Grant No. 813327) and La Ligue Nationale contre le Cancer (Grant No. TDLM23697). M.A.M-R. acknowledges support by the Spanish Ministerio de Ciencia e Innovación (PID2020-115696RB-I00 and PID2023-151484NB-I00).

## Notes

### Competing Interest Statement

The authors have declared no competing interest.

